# A mechanism for the proliferative control of tissue mechanics in the absence of growth

**DOI:** 10.1101/266577

**Authors:** Min Wu, Madhav Mani

**Author notes:** /.

## Abstract

During the development of a multicellular organism, cells coordinate their activities to generate mechanical forces, which in turn drives tissue deformation and eventually defines the shape of the adult tissue. Broadly speaking, it is recognized that mechanical forces can be generated through differential growth and the activity of the cytoskeleton. Based on quantitative analyses of live imaging of the *Drosophila* dorsal thorax, we suggest a novel mechanism that can generate contractile forces within the plane of an epithelia - via cell proliferation in the absence of growth. Utilizing force inference techniques, we demonstrate that it is not the gradient of junction tension but the divergence of junction-tension associated stresses that induces the area constriction of the proliferating tissue. Using the vertex model simulations, we show that the local averaged stresses can be roughly elevated by a fold of *p* 2 per cell division without growth. Moreover, this mechanism is robust to disordered cell shapes and the division anisotropy, but can be dominated by growth. In competition with growth, we identify the parameter regime where this mechanism is effective and suggest experiments to test this new mechanism.

## Introduction

During morphogenesis an organism grows in size and undergoes successive deformations at the cellular and tissue-wide scales. Driving these changes are the cellular processes of growth and cytoskeletal activity [1]. The coordination of these processes can generate long-range forces and thus can change the size and shape of tissues, organs, and the whole organism. For epithelial morphogenesis, cell growth and divisions, regulated by growth factors and mitogens [2], can expand the tissue area and can induce non-trivial planar stress patterns. For example, fast growing clones can generate tension in the surrounding slower growing clones [3–5]. Besides differential growth, tension can be generated by actin-based contractility mediated by motor proteins. The enrichment of the pool of molecular motors on the apical surface of the cells has been show to play a role in the constriction of the apical surface of cells [6–8], while the enrichment of molecular motors on a single cell-cell junction plays a role in the constriction of the junction’s length thus cell-cell intercalations [9–12]. An aggregate of such cellular effects can generate large scale contractile forces and deformations. Taking *Drosophila* gastrulation, for example, a gradient of motor protein activity drives tissue flow from lower- to higher-concentration regions of motor proteins [6, 7]. Thus, an understanding of morphogenesis would involve not just a list of the patterning factors, but also an understanding of how the patterning factors impinge on cellular mechanisms to give rise to biological form. In particular, how do cellular activities affected by the role of regulatory factors – such as morphogens and stresses –pattern and update internal stresses and thus give rise to tissue deformations? Furthermore, do cellular activities generate forces in other ways?

In this paper, we propose that in the absence of cell growth, cell division is an alternate mechanism –alternate to the gradient of motor protein activity–to generate constricting forces during tissue morphogenesis. Our proposal is based on our analyses of live-imaging data of the epithelium morphogenesis in the *Drosophila* dorsal thorax (notum) during metamorphosis [13, 14]. Although cell division is coordinated with growth in most tissues, it is not the case in the notum starting from 11 to 35 hours after pupa formation - the epithelial tissue does not increase in its overall size while the total cell number more than doubles [13, 14]. In the most active tissue region of the notum, we identify a time window when the tissue reduces its size while cells divide. By using an inference scheme [15], we exclude the possibility that the tissue constriction is due to the elevated junction tension, which would suggest a higher level of motor protein activity per junction. By using a vertex model, we confirm that a higher level of tensile stresses can be generated by division in the absence of growth alone. We then find the signature of this simple mechanism in the notum data. Finally we identify the parameter regime where this mechanism is effective. Theoretical considerations are summarized in *Materials and Methods*. In *Discussion*, we summarize our findings and suggest experiments to test the newly proposed mechanism.

## Materials and Methods

### The vertex modeling of epithelial tissue mechanics

For simulation results in figure 3, 4 and 6, we adopt the vertex-based modeling [16] to describe tissue dynamics. Vertex models have been used to study many epithelial systems, such as in the *Drosophila* embryo [17], wing disc [4, 18, 19], pupa abdomen [20] and *Xenopus* embryo [21, 22]. See recent papers [23–25] for more details in the numerical implementation [23, 24] and their applications in studying biological systems [25]. In the vertex model, the activities of cells - divisions, the rearrangement between adjacent cells, as well as the delamination of cells from the epithelia are all captured by the addition and deletions of junctions and vertices from the lattices. Cell sizes and shapes are resulted from positions of the vertices, updated according to the following constitutive equations. On each vertex *i*, the update of its position is according to the force balance

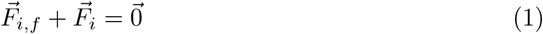

between the friction forces 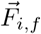 and the internal forces of the epithelium 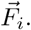 For the friction forces, we consider

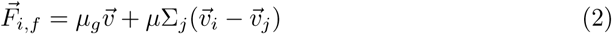

both from the substrate by a constant friction coefficient *μ_g_* and from the neighboring materials by a constant friction coefficient *μ*. *j* enumerates the neighboring vertices connecting to the vertex *i*. The internal force 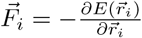 is derived from the energy functional:

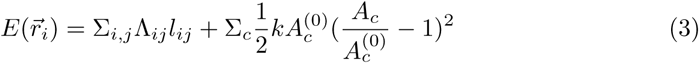

where *Λ_ij_* is the junctional force, considered to be uniform in our study, set to be a constant Λ, and *l_ij_* is the length of the junction connected from vertex *i* to vertex *j*. The second term considers the penalty of cell apical surface area *A_c_* deviated from the target area 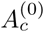 by a constant *k* as the bulk modulus. *c* enumerates the neighboring cells in contact with the vertex *i*. By prescribing 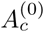 and *k* for individual cells as well as Λ for all the junctions, we simulate the tissue configuration by solving the vertex system to mechanical equilibrium. A division is modeled by adding a new junction on a polygonal cell with a random angle orientation, which divides the mother polygonal cell into two daughter polygonal cells. The growth per division is modeled by a single parameter α, defined as the area ratio between two daughter cells (each with area 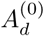 and the mother cell (with area 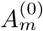), satisfying 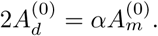 When *α* = 1, cells only divide without growth. 1 < *α* ≤ 2 corresponds to the cell division where the growth is also taking place. *α* = 2 corresponds to the situation where division and growth are perfectly coordinated – each daughter cell has exactly the same target size as the mother cell. *α* > 2 corresponds to that the growth rate is faster than the division rate, which is beyond the scope of this study. For rearrangement, one pre-existing junction between two previously connected cells disappears and a new junction between the newly connected cells appears (called T1 transitions). When one junction *l_o_* goes below a user-specified threshold *T_h_*, based on the current configuration involving the four cells surrounding the junction *l_o_*, we generate an alternative configuration, where the old junction *l_o_* is rotated by *π*/2 with the same length. We compare the total energy of the current and alternative configurations involving the four cells and pick the configuration with lower total energy. For delamination, we extrude a cell when its edge number is below 4 (called T2 transitions). In this study, since the focus is the effect of division without growth, for most simulation results we consider *α* = 1 (Figure 3 and 4). The exception is figure 6 where we explore the parameter space 1 ≤ *α* ≤ 2 while changing the non-dimensional parameter 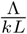 (*L* is the characteristic length, taken to be the junction length of the original hexagonal tissue lattice before cell divisions).

### The stress tensor from the vertex model

The averaged tissue stresses in Figure 3 and 4 are computed by the following two steps. At first, we compute the average stress tensor of individual cells by integrating the external traction along the cell boundary averaged by the cell area 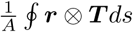 [26, 27]. This gives the stress tensor of the cell

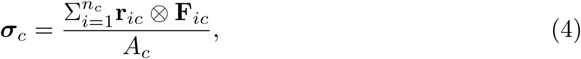

where r_*ic*_ is the position on the vertex *ic* and F_*ic*_ is junctional force at vertex *ic* from the junction connecting to the external vertex. Notice this equation is equivalent to the equation to compute the stress tensors in [28]. Then we measure the average tissue stress tensor by

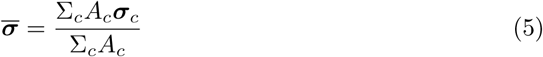

where *A_c_* is the area of the cells and *σ_c_* is the cell stress. The average tissue stress component *σ* is computed by 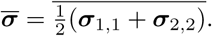

## Results

### The size reduction of the dividing tissue

Analysis of the live-imaging (kindly provided by the Bellaïche lab) taken of the *Drosophila* notum (Figure 1a) demonstrates that higher proliferating regions of the tissue have higher rates of constriction. In particular, tissue regions (patches from the top two rows in figure 1b-e) distant from the midline have an inverse correlation between profiliferation and constriction rates during the period of ∼ 17 *−* 21 hours after pupa formation. This seems to run counter to the differential growth hypothesis, wherein cell proliferation is associated with tissue growth and local dilation [3, 4, 29]. Our hypothesis is that in the absence of growth, the introduction of a new junction following a cell-division [30] will result in a new source of tensile stress, which can lead to apical constriction (Figure 2a).

**Fig 1.**
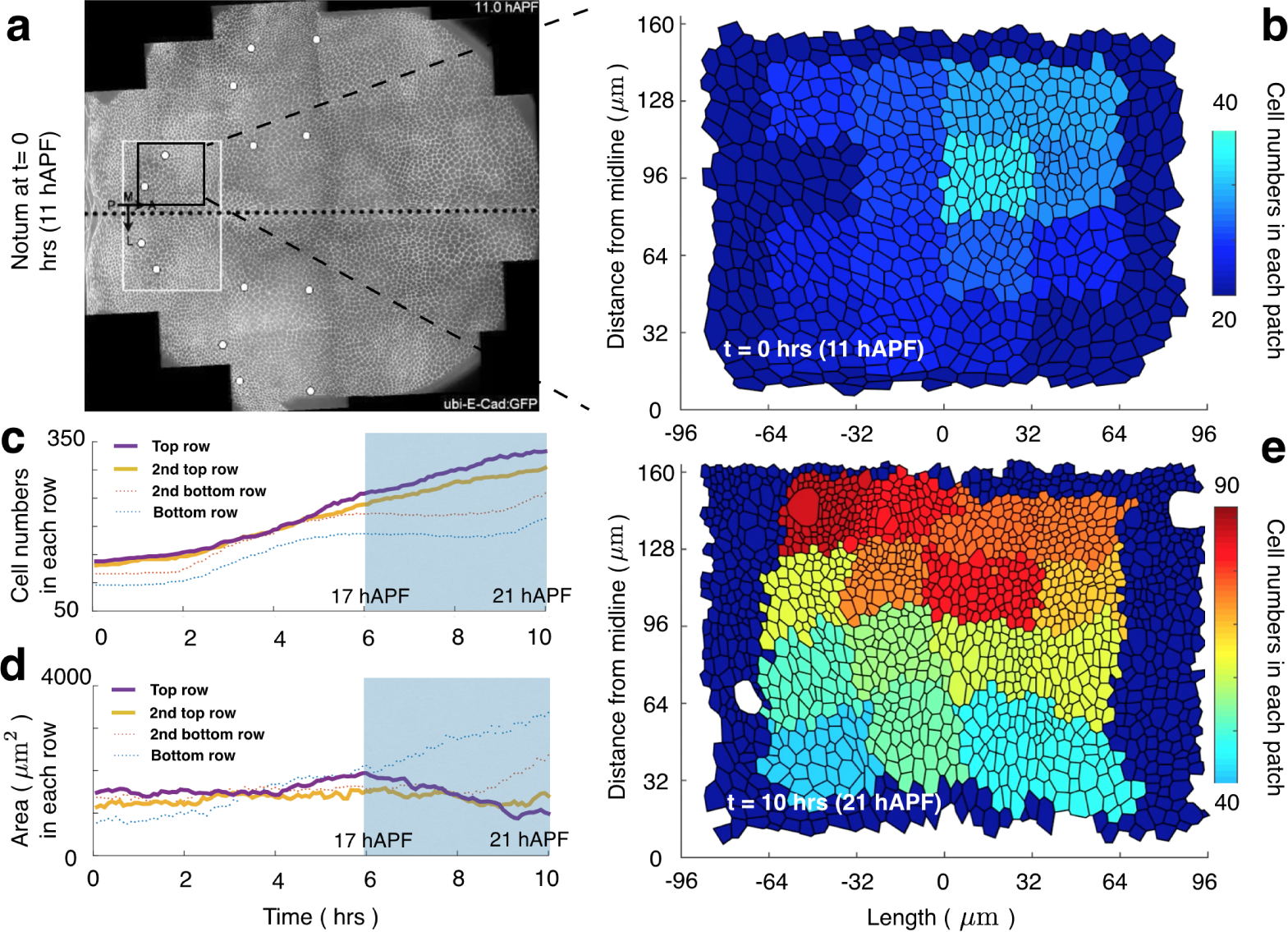
Quantification of cell numbers and tissue size from data. **a** shows the notum at time *t* = 0, 11 hours after pupa formation (hAPF). The tissue in the black squared region is divided into 4 *×* 4 equal area patches and the color presents the number of cells in each patch. **b** shows the patches at *t* = 0. The patches are tracked (see method) and e shows the deformed 4 *×* 4 patches at *t* = 10. Notice the top row tissues are relatively smaller than the bottom row tissues. The total cell number and the area of each row versus time are quantified in **c** and **d**, respectively.

**Fig 2.**
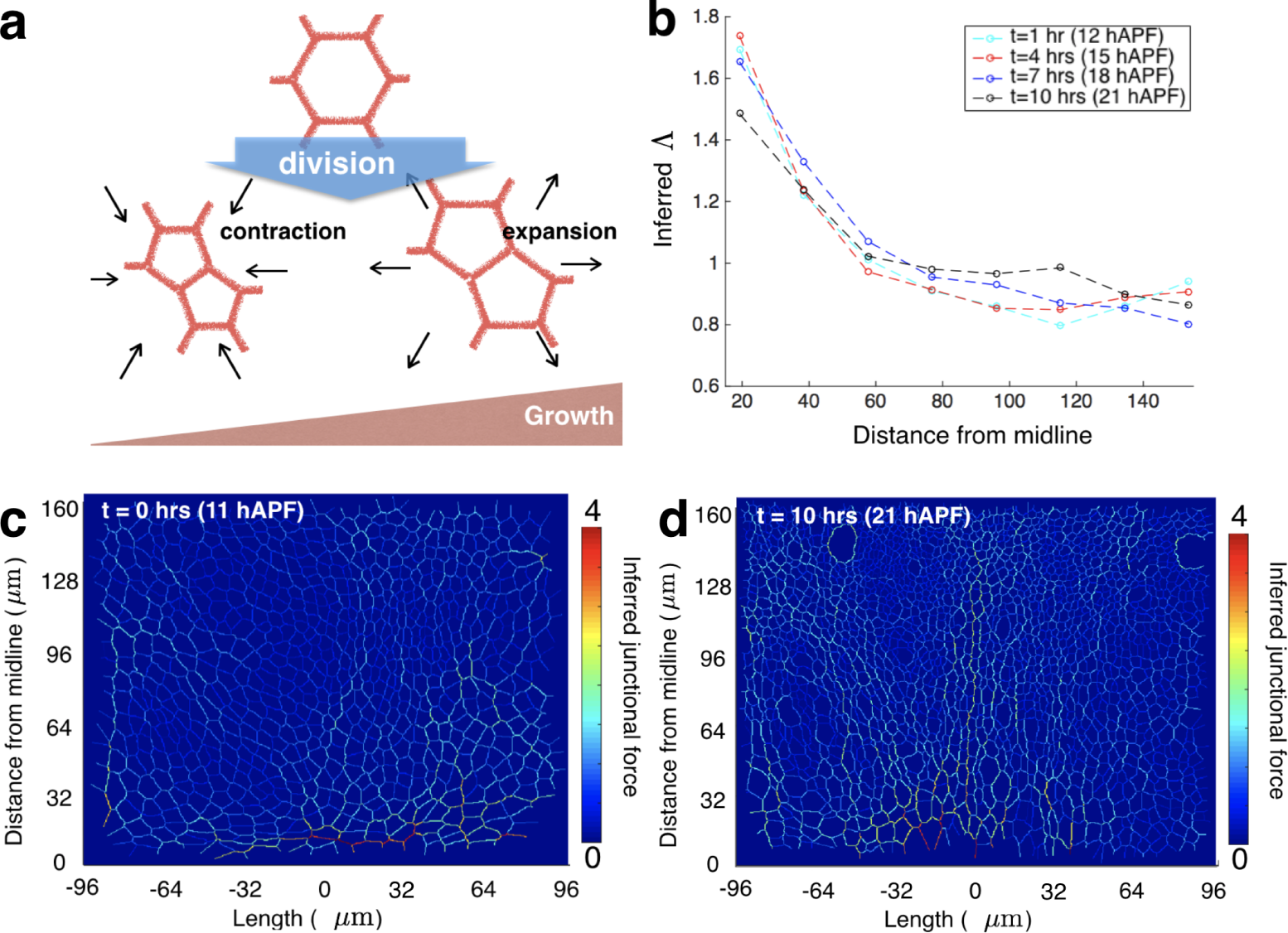
Schematics and junction tension inference. **a** shows the total size of apical surface area of the two daughter cells is modulated by growth. **b** plots the inferred junction tension versus the distance from the midline. Cell-cell junctions are sorted into 8 bins according to their distance from the midline and the average tension is calculated over each bin. c and d show the heat map of the relative junction tension at *t* = 0 and *t* = 10, respectively.

### The inference of junction tension distribution

Is there an evidence for the aforementioned hypothetical mechanism being responsible for the observed constriction in the data (Figure 1d-e)? A natural alternative hypothesis is that the size reduction of the proliferating tissue may be a result of elevated tension along junctions. In the absence of being able to experimentally measure the membrane-tensions in cells in the different regions of the tissue, we rely on force inference techniques [15] that give us access to the relative spatial distribution of tensions. Our results indicate that the average tension is not elevated in the constricting region distant from the midline (Figure 2b-d). In detail, we infer the junction tension distribution at different time points from 11 to 21 hours after pupa formation (hAPF). For illustration, Figure 2c (d) shows the heat map of junction tension at 11 (21) hAPF, denoted as *t* = 0 (*t* = 10). At different time points (see figure 2b), we plot the averaged inferred junction tension versus the distance from the midline. To do this, cell-cell junctions are sorted into 8 bins according to their distance from the midline and the average tension is calculated over each bin (8 data points on each dashed curves in figure 2b). Notice the data points with distance *>* 80 microns corresponds to the constricting region. One can see that the relative distribution of average junction tension along the distance from the midline does not change significantly over time, suggesting the tissue constriction in the faster proliferation regions is not due to the elevation of junction tension. We thus anticipate that the tissue constriction is due to the addition of new junctions that contribute to the tensile stress components within the epithelial plane.

### The elevation of tissue tensile stresses due to divisions

In order to show how much extra planar stress a single cell-division event contributes, we first focus on a simple situation. We consider regular polygonal cells that have a size *R*_0_ and have a uniform junction tension, Λ. The size *R*_0_ is defined as the radius of the circle inscribing the polygon, and there are zero stresses from the cell interior. For all the polygonal cells with different number of sides, we find their stress tensors to be isotropic (see Eq. (4)). In figure 3a, we show that the single diagonal stress component, *σ*, of the regular polygonal cells scales with Λ/*R*_0_. While the proportionality between *σ* and Λ/*R*_0_ depends on *n*, the number of sides a cell has, figure 3a demonstrates that this proportionality does not vary much between *n* = 5 to 7, which is the case for most cells. If we assume cells remain as regular polygons, division without growth will reduce its size 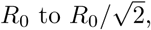 then will increase the stress by a factor of 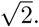 Without assuming cells are regular polygons, we implement the vertex model summarized in *Materials and Methods* to show that the predicted proportionality between *σ* and Λ/*R*_0_ still holds with disordered cellular lattices. Starting with a tissue tiled by regular hexagonal cells with areas equal to the targeted area (see more details in *Materials and Methods*), we perform two rounds of divisions of cells with random division orientations with no growth (See figure 3c-d for the simulated tissue topography). In both rounds of divisions, the average stress tensors, calculated from Eq. (4) and (5), maintain their isotropic nature, and the diagonal component *σ* scales with Λ/*R*_0_. This result agrees with our prediction of stresses calculated by assuming all cells are regular polygons with the same area as their irregular counterparts in the simulation (See figure 3b). In addition, the ratio of *σ* between the two successive division rounds is approximately 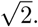 Altogether, these findings show that the generation of tensile stresses per round of cell divisions in the absence of growth is 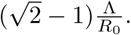

**Fig 3.**
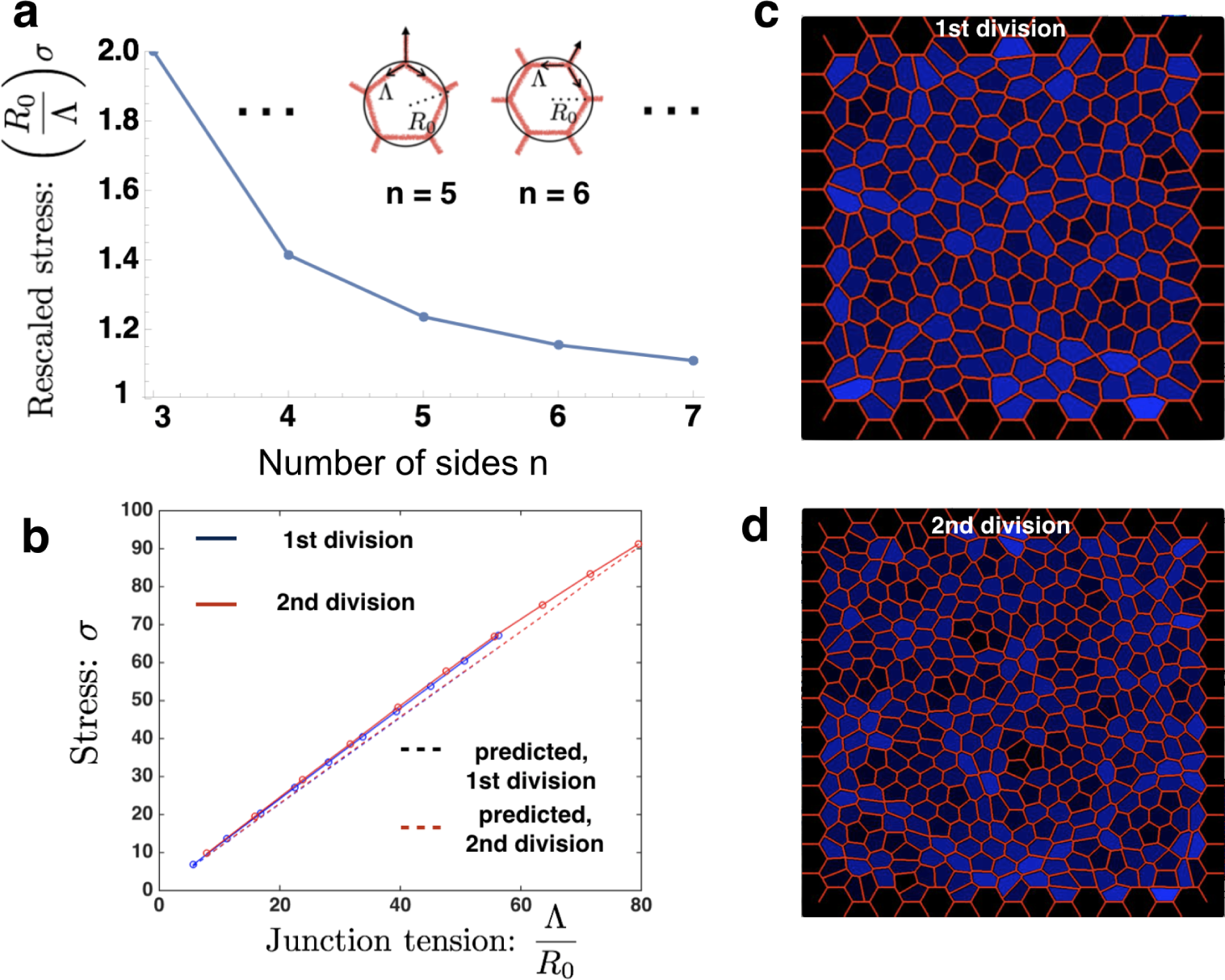
Stress analysis and dividing tissue area changes. **a** shows stresses of regular polygons rescaled by Λ/*R*_0_ versus the number of sizes n. **b** shows the isotropic component *σ* of the averaged stress over the computational also scales with Λ/*R*_0_ (solid lines) and agrees well with the stress components calculated by fitting all cells as regular polygonal cells with equal sizes (dashed lines). c and d show the tissue overlay after first and second rounds of divisions with random division orientation on a hexagonal lattice. The color gradient from black to blue on each cell color-codes its isotropic stress component *σ* relative to the average.

### Biased cell division orientation

So far, we have shown the tissue stress can be elevated due to division without growth, because the stress component *σ* is inversely proportional to the cell size *R*_0_. This result is based on the implementation of Eq. (4) and (5) on regular polygonal cells and tissues with random division. In the case of random divisions, such as in figure 3c-d, the stress tensor is approximately isotropic. In this section, we ask if biased divisions in a preferred direction in a patch of tissue would alter our conclusion. Based on the tissue overlay in figure 3c, we investigate the effect of anisotropic cell divisions by introducing new junctions with their angles drawn from the uniform distribution [*π*/2 − *π* (1 − *β*), *π*/2 + *π* (1 − *β*)], *β* being the level of division anisotropy. See figure 4a for simulated tissue topography with *β* = 0, 0.5 and 1. As *β* increases, the stress tensor becomes more anisotropic. In figure 4b, we plot the stress component *σ*_‖_ (*σ_⊥_*) in the direction aligned with (orthogonal to) the mean division axis, varying *β*. Not surprisingly, *σ*_‖_ is larger than *σ*_⊥_, and the stress anisotropy is pronounced as *β* increases. Interestingly, the isotropic component of the stress tensor, 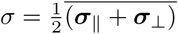 is still 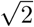 fold of that before division (Figure 4b, the two black dashed lines). This can be explained by the average tissue stress equation

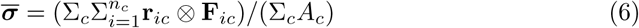

derived from Eq. (4) and Eq. (5), which gives 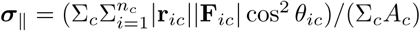 and 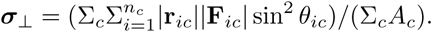 *θ_ic_*’s are the angle between each junction and the mean division axis. 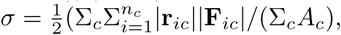 and is independent of the angles *θ_ic_*’s. We conclude that the isotropic stress component *σ* is not affected by the division orientation bias, and the tissue constriction is guaranteed when the fixed boundary is relaxed.

**Fig 4.**
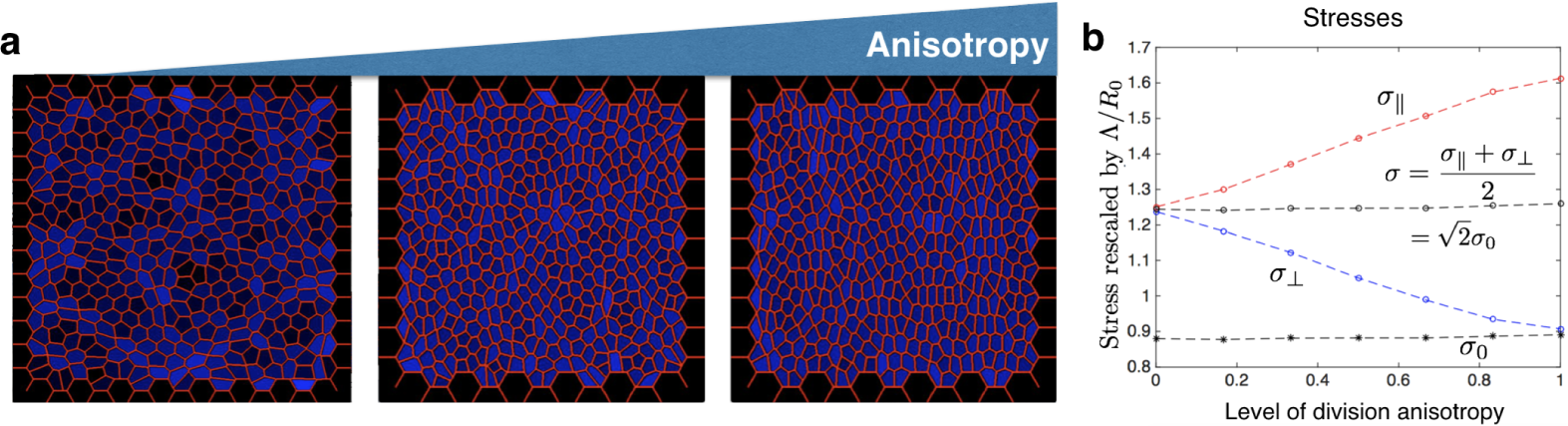
Stress modulated by division anisotropy. **a** shows divided tissue patches with fixed boundary resulted from different level of anisotropy of division orientation. The left figure shows the random oriented division, the middle figure shows the division orientation drawn from [*π*/4, 3 *π*/4] and the right figure shows the division orientation drawn from [0, *π*]. The color gradient from black to blue on each cell color-codes its isotropic stress component *σ* relative to the average. **b** shows the stresses rescaled by Λ/*R*_0_, where *R*_0_ is the average cell size after division. *σ*_‖_(*σ*_⊥_) shows the stress in parallel with (orthogonal to) the mean division axis *θ* = *π*/2. *σ*_0_ denotes the original average stress before division.

### The signature of the new mechanism in the data

Thus far, we have shown that in theory cell divisions can increase the tissue tensile stress up to 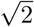 of the original stress magnitude. Notice this analysis is done with fixed boundary conditions, thus we are estimating the “net” tensile stress added into the tissue system before deformation. The deformation of the tissue patch in the notum data may be driven by other sources of forces besides the junction tension. In particular, the constriction of the proliferating tissue can be a result of 1) the relaxation of the tensile forces from the upper boundary of the tissue patch; and/or 2) other sources of stress divergence. However, it is not likely that the tension from the upper boundary is relaxed since the tissue patch is advancing upward (data not shown). If the increase of junction-tension associated stresses due to division is the driver of the observed constriction, one expects to see an increasing gradient of Λ/*R*_0_ from the midline to the distant constricting region. To quantify the spatial distribution of Λ/*R*_0_ (see figure 5b), we further calculate the averaged cell *R*_0_ from the midline (see figure 5a), and calculate Λ/*R*_0_ by dividing the averaged (plotted in figure 2b) by the averaged *R*_0_. Indeed, Λ/*R*_0_ in the distant regions elevates (black dashed line in figure 5b) after the cells have divided - in agreement with our argument that the increase of junction-tension associated stresses due to division in the absence of growth is the major cause of the observed constriction.

**Fig 5.**
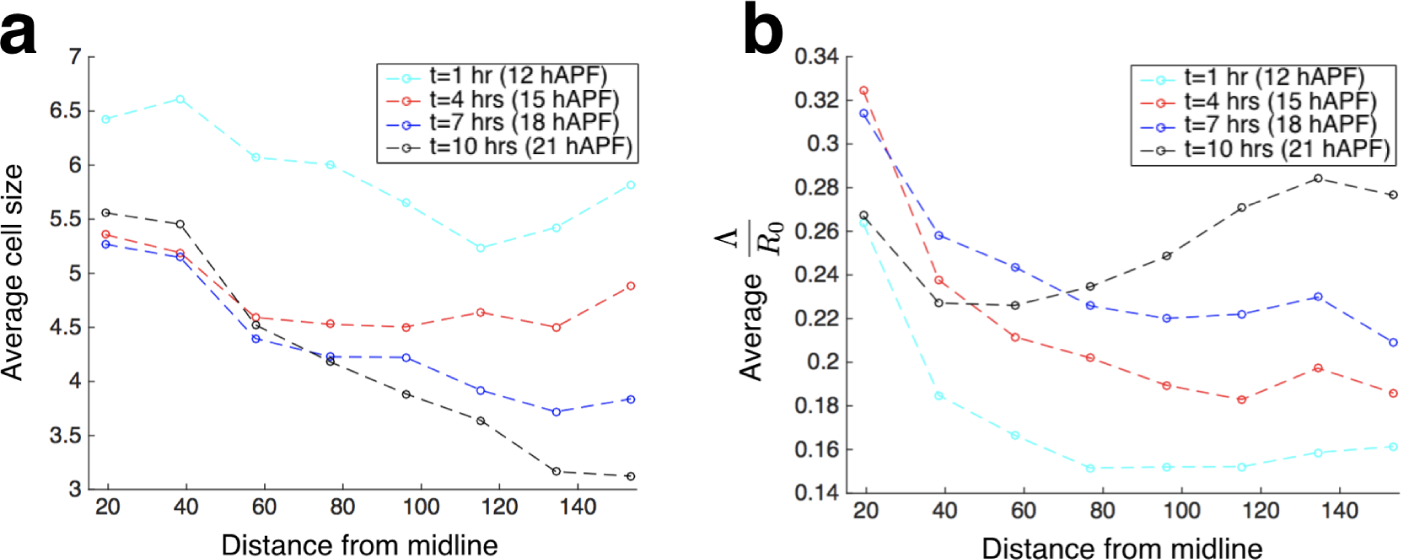
The planar cell size *R*_0_ and the scale of the stress Λ/*R*_0_ from data. **a** plots the cell size *R*_0_ versus the distance from the midline. *R*_0_ is calculated as the radius of the circle inscribing a regular polygon with equal area. Cells are sorted into 8 bins according to their distance from the midline and the average *R*_0_ is calculated over each bin. **b** plots the average Λ/*R*_0_ versus the distance from the midline.

### The competition between growth and junction tension

The elevation of junction-tension associated tensile stress induced by division should be inherent in proliferating tissue regions. However, more frequently proliferation is associated with both growth and division, where in total the constricting trend of the dividing tissue is dominated by tissue expansion induced by growth. We can think of the growth-induced area expansion as caused by a cell internal pressure due to the mismatch between the current planar cell area *A* and the target cell area *A* ^(0)^. In the competition between the two opposing effects, we use the vertex model to identify the condition under which the constricting regime emerges. In the vertex model, the cell internal pressure *P* = *k* (1 − *A/A*^(0)^), proportional to a parameter defined as the bulk modulus *k* (see the *Materials and Methods*). To quantify the relative growth rate over division rate, we introduce a single parameter *α*, defined as the area ratio between two daughter cells (each with area 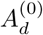) and the mother cell (with area 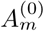), satisfying 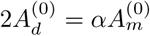 (see more details in *Materials and Methods*). When *α* = 1 cells divide without growth, while when *α* = 2 the cell-division rate perfectly coordinates with the growth rate– giving two daughter cells exactly the same size as the mother cell. We simulate a patch of proliferating tissue under free boundary conditions (without considering the stretching or compression by external forces), so the tissue-area changes are solely tuned by the competing effects between the introduced compression from growth and the introduced tension from division. Under this condition, we quantify the tissue size variation – the ratio between the area of the tissue after division and before division– by changing *α* and the dimensionless parameter 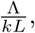 and plot the phase diagram in figure 6. Here, *L* is the characteristic length, taken to be the junction length of the original hexagonal tiling of the tissue before cell divisions. The tissue area increases after cells divide, when 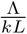 is small or *α* is closer to 2; while the tissue area decreases only when 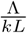 is large and *α* is closer to 1. Notice when 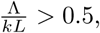 many cells collapse into points, reminiscent of the delamination of epithelial cells. The outcome of delamination has also been found by previous theoretical works [18, 21, 22], and its effect on tissue constriction is beyond the scope of this study. Considering the parameter space where delamination does not occur and the growth rate at most matches the division rate (*α* ≤ 2), we show that the proliferative tissue constricts less frequently than it expands, from a probabilistic point of view.

**Fig 6.**
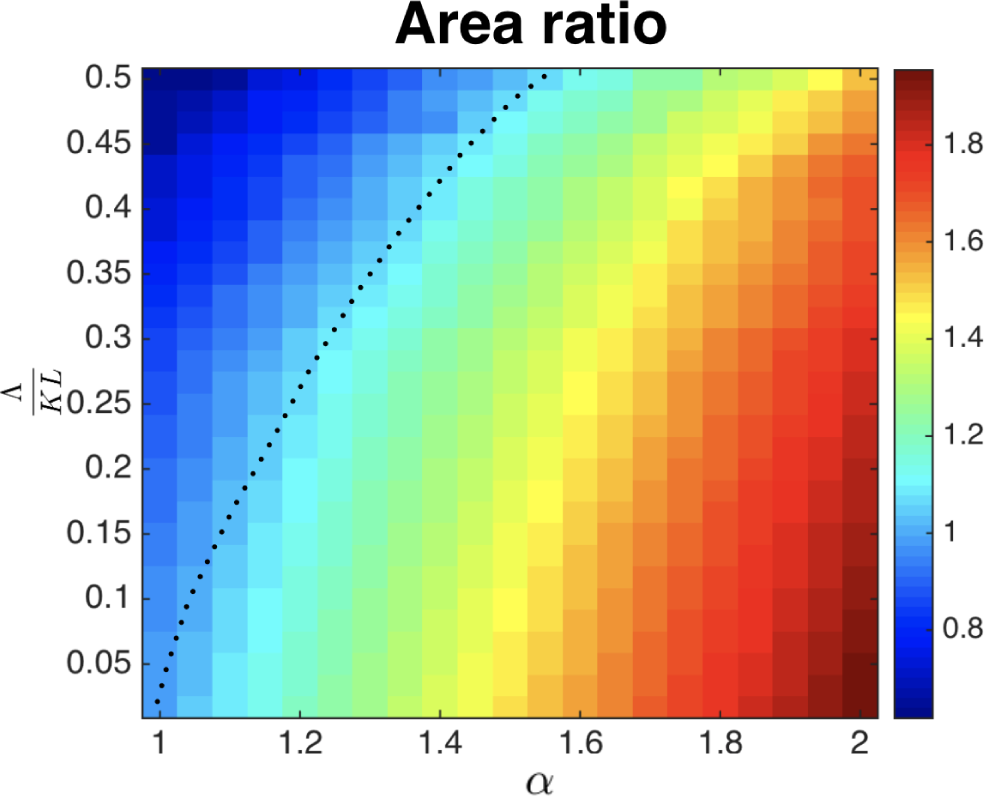
The ratio between the area of the tissue after division and before division. The area ratio is modulated by the growth parameter *α* and the dimensionless parameter 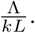

## Discussion

In this paper, we propose a novel mechanism that may play a role in the patterning of forces in epithelial tissues - the generation of contractile forces due to proliferation in the absence of growth. Focusing on the region of the tissue that are most proliferative (Figure 1), we identify a time window during which cells proliferate while the apical surface area of the proliferative tissue decreases (Figure 1c-d). It is plausible that the creation of a junction that separates two daughter cells in the absence of growth can produce a contractile force locally, and that aggregates of cell divisions without growth can work as an alternative mechanism for tissue constriction, besides tissue-wide motor protein gradient. It has been shown earlier in the *Drosophila* larval wing disc that the regulation of cell (mass) growth is independent of the cell cycle regulations [31–33], which suggests that growth rates and division rates can be regulated separately. In theory, we show that without considering growth, division alone can elevate the planar tensile stress up to 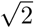 fold of its original magnitude, and the tensile stress is generated because the stress components scale inversely proportional to the planar size of the epithelial cells *R*_0_ (see Figure 3 and the results above). We further show that this constricting mechanism works even when the orientation bias is given to cell divisions (Figure 4). Using the notum data, we have shown that the tension associated stress component is relatively elevated in the constricting region (Figure 5). Lastly, we have shown that proliferation induced tissue constriction is not likely to happen when there is moderate growth or low averaged junction tension (Figure 5). From a probabilistic point of view, proliferation induced tissue size reduction occurs less frequently than tissue expansion.

In the data, most cells divide twice (Figure 1 c), while the overall tissue does not grow in size. Interestingly, we only see the division with tissue constriction in the second round of division while in the first round, their sizes stay nearly the same (top two rows in Figure 1 b-e). Our theory in this paper suggests that the generation of tensile stresses per round of cell divisions in the absence of growth is 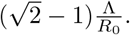 Thus, the magnitude of the stress generation is determined by 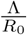 initially. When Λ is small, or *R*_0_ is large, the increase of stress due to division may be negligible. It has been reported that both the junction tension, Λ, and the level of myosin II activity along the junction in the notum increase with the pupal age [34]. In addition, the averaged cell size *R*_0_ in the second round of division is smaller compared to the averaged cell size *R*_0_of the first round. Furthermore, during the first round of cell division, there is a considerable amount of myosin generating constricting forces from the medial pool of the cells [34], which can counterbalance the increase of stress due to division. These altogether may explain why we only see the correlation between the tissue constriction and cell-number increase in the second round of division.

It has been shown earlier in [14] that the cell divisions are linked to the epithelial mechanics in the notum. When cell divisions are blocked, both the pattern of stresses and the contraction elongation at the tissue level are disrupted compared to the wildtype. Given the spatiotemporal heterogeneous patterning of cell proliferative events and division orientation, and the uncertainty of the contribution from growth per se, it is hard to draw the conclusion that division has resulted in tissue constriction. Based on our theoretical studies, we suggest that future studies increase the junctional tension on the entire notum and quantify if the correlation between division and area reduction is accentuated during the constricting phase. To do this, one could increase the level of myosin II activity via perturbing the function of the Myosin activator Rho kinase (Rok) [35, 36], which has been successfully implemented in [34].

